# Influenza virus inhibits respiratory syncytial virus (RSV) infection via a two-wave expression of interferon-induced protein with tetratricopeptide (IFIT) proteins

**DOI:** 10.1101/2020.08.17.253708

**Authors:** Yaron Drori, Jasmine Jacob-Hirsch, Rakefet Pando, Aharona Glatman-Freedman, Nehemya Friedman, Ella Mendelson, Michal Mandelboim

**Author notes:** **Corresponding authors**: Michal Mandelboim. **Declaration of interest**: All authors declare no conflict of interest.

## Abstract

Influenza viruses and respiratory syncytial virus (RSV) are respiratory viruses that primarily circulate worldwide during the autumn and winter seasons. Seasonal surveillance shows that RSV infection generally precedes influenza. However, in the last four winter seasons (2016-2020) an overlap of the morbidity peaks of both viruses was observed in Israel, and was paralleled by significantly lower RSV infection rates. To investigate whether the influenza virus inhibits RSV we performed coinfection of Human cervical carcinoma (HEp2) cells or mice with influenza and RSV and we observed that the influenza inhibited RSV growth, both in vitro and in vivo. Mass spectrometry analysis of mouse lungs infected with influenza identified a two-wave pattern of protein expression upregulation, which included members of the interferon-induced protein with tetratricopeptide (IFITs) family. Interestingly, in the second peak of upregulation, influenza viruses were no longer detectable in mouse lungs. We also observed that knockdown and overexpression of IFITs in HEp2 cells affected RSV multiplicity. In conclusion, influenza infection inhibits RSV infectivity via upregulation of IFIT proteins in a two-wave modality. Understanding of the interaction between influenza and RSV viruses and immune system involvement will contribute to the development and optimization of future treatment strategies against these viruses.

**Author Summary:** Respiratory syncytial virus (RSV) and influenza viruses are both respiratory viruses associated with morbidity and mortality worldwide. RSV is usually detected in October, with a clear peak in December, whereas influenza virus arrives in November and peaks in January. In the last four seasons, influenza infection overlapped with that of RSV in Israel, which resulted in decreased morbidity of RSV suggesting that influenza virus inhibits RSV infection. To identify the mechanism responsible for the influenza inhibition of RSV we performed experiments in culture and in mice. We observed that influenza infection results in two wave modality of inhibition of RSV infection. Using mass spectrometry perfornmed on lungs from infected mice we show that influenza infection induces the expression of (IFIT) family of proteins which also showed a two-wave modality. Using knockdown and overexpression experiments we showed that indeed the IFTIs inhibits RSV infection. Our study provides new insights on the interaction between influenza and RSV viruses and immune system involvement and contribute to the development of future treatment strategies against these viruses.

## Introduction

Respiratory syncytial virus (RSV) and influenza viruses are both respiratory viruses associated with significant morbidity and mortality worldwide(1). RSV is a leading pathogen causing acute lower respiratory tract infection (ALRTI)(2) in all age groups(3, 4) primarily infants and young toddlers(5, 6), while influenza virus affect all age groups(7). In Israel, RSV is usually detected in October, with a clear peak in December, whereas influenza virus arrives in November and peaks in January(8). Previous surveillance demonstrated a close temporal relationship between circulating influenza virus and RSV, where influenza epidemics usually occur when RSV infections subside(9). On the other hand, in the last four seasons, influenza infection overlapped with that of RSV in Israel, which resulted in decreased morbidity of RSV suggesting cross-immunity between influenza virus and RSV infection(10). We assume that changes in the arrival time of one virus can affect the arrival time and morbidity pattern of the other.

The activity of both innate and adaptive immune systems against both viruses is very well documented(11, 2). Innate immunity against these viruses includes, among others, interferon stimulating genes (ISGs) pathway, which trigger the expression of the interferon-induced protein with tetratricopeptide (IFIT) family of proteins (12, 13). In humans, IFIT1-3 proteins form a complex with each other to inhibit translation of viral mRNA molecules(14). IFIT1 acts as a sensor that identifies specific viral single-stranded RNAs and selectively inhibits viral protein synthesis without affecting host cell protein synthesis(15). While IFIT1 is known to specifically recognize the viral mRNA(15), it has been suggested that IFIT2 and IFIT3 facilitate the binding of the IFIT1:2:3 complex structure to the viral mRNA(17, 16). Less in known about the function of IFI44. The present work investigated the role of IFIT1-3 and IFI44 proteins in the cross-immunity effect observed between influenza and RSV.

## Results

### Influenza infection inhibits RSV

Samples obtained from both hospitalized and non-hospitalized patients in Israel 2009-10, showed that influenza virus A/H1N1pdm09 virus that year was present in the country during the summer season and peaked in July and November 2009 (Fig 1A). In contrast, in winter seasons of 2013-16, RSV infection was detected in October, when it usually appears and peaked in December and November(18), while influenza infections peaked in February (Fig 1B) and January (Fig 1C). Surprisingly, from the winter of 2016-17 onwards, both RSV and influenza co-circulated in October and both peaked around January (Fig 1B and C). this early circulation of influenza in the winters of 2016-20 coincided with a lower morbidity of RSV than previous seasons (2013-16), and also coincided with later peak of RSV in January. Overall, RSV morbidity percentages were significantly lower (p<0.0005) as compared to the two preceding winter seasons (Fig 1B and 1C), suggesting that influenza infection affecting RSV infection rate.

**Fig 1.**
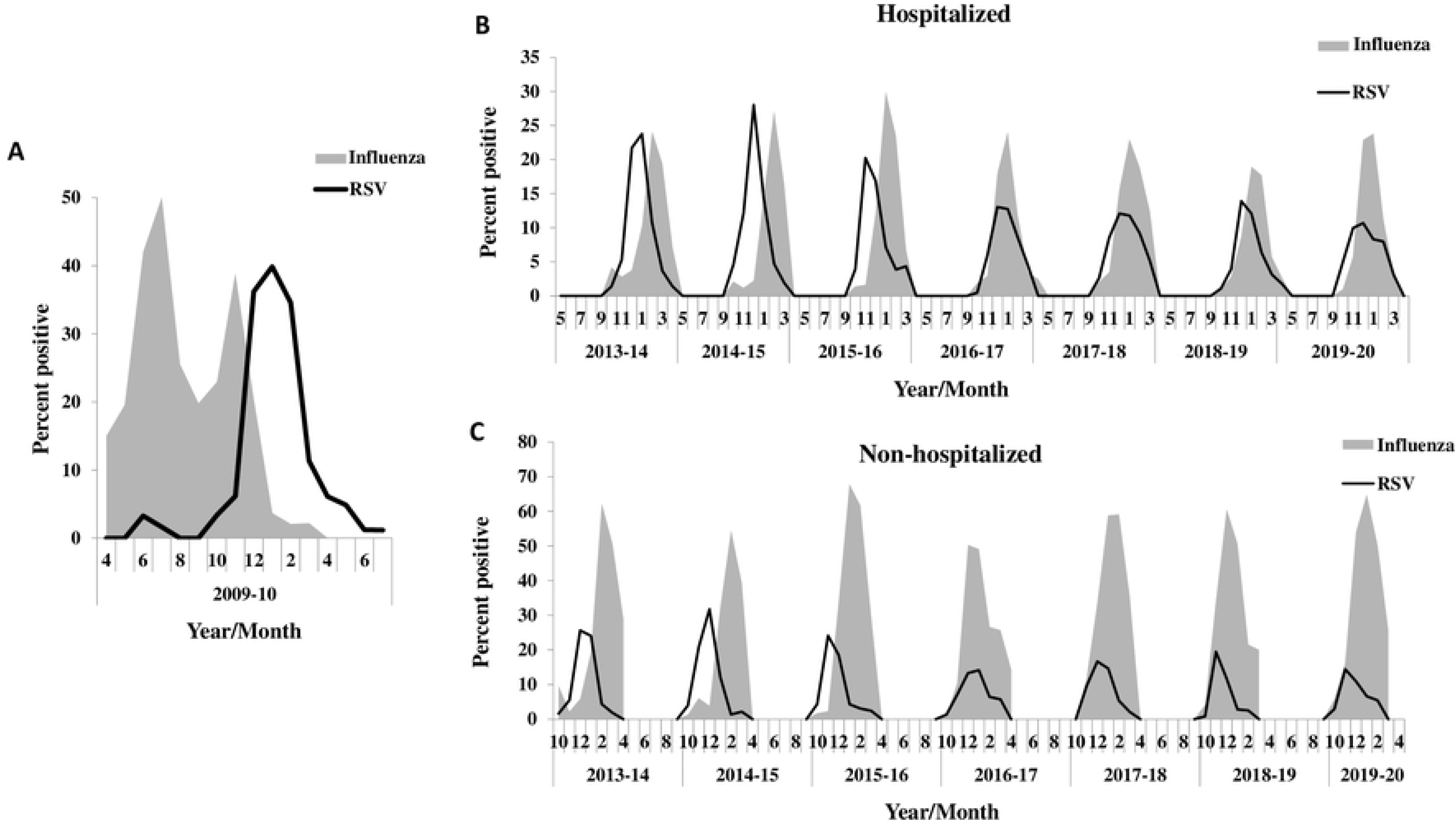
An overview on the percentage of influenza-positive versus RSV-positive samples in the winter seasons 2013-2020. **(A)** The weekly percentage of positive cases of infection with respiratory viruses (RSV, and/or influenza) from 2009-10. **(B)**. The weekly percentage of positive cases of infection with respiratory viruses (RSV, and/or influenza) from 2013 to 2020 among hospitalized patients. **(C)** The weekly percentage of positive cases of infection with respiratory viruses (RSV, and/or influenza) from 2013 to 2020 among non-hospitalized patients. The black line represents the infection patterns for RSV and the gray area represents influenza infection patterns. The percentage of patients bearing each virus was calculated each month in relation to the total number of samples tested for the presence of that virus.

### Influenza infection inhibits RSV in vitro

To test whether influenza virus presence can indeed inhibit RSV infection rate, HEp2 cells were infected with RSV or with influenza A virus, followed by RSV; the number of RSV copies was then determined at various time points. When applied alone, RSV copy numbers increased with time from infection (24, 48, 72, 96 and 144 hours) (Fig 2), however, when first infected with influenza A, cells produced statistically significant fewer RSV copy numbers at all tested time points (Fig 2).

**Fig 2.**
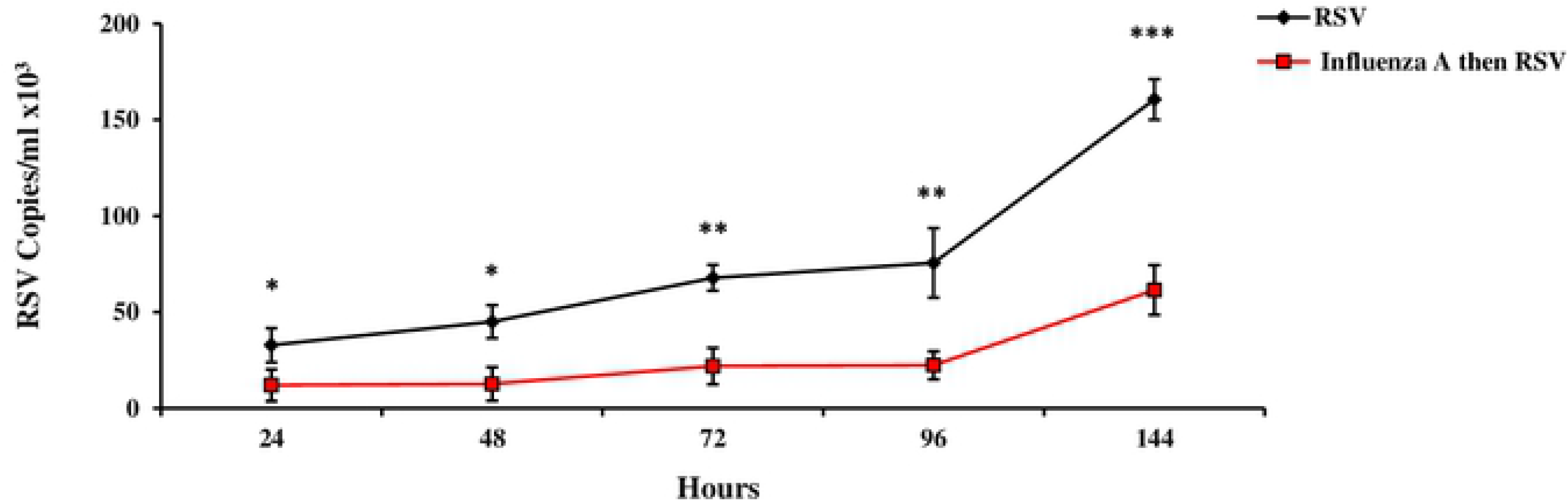
Co-infection of HEp2 cells with RSV and influenza A/H3N2. HEp2 cells (1×10^6^) were infected with RSV (6×10^5^ PFU, 3 h), and then with influenza A/H3N2 (6×10^5^ PFU, 3 h). Non-infected cells served as control. RNA was extracted from supernatant samples collected at predefined time points (24-144 h). qRT-PCR was performed to test for viral quantity/presence. The black line indicates infection with RSV, the red line indicates infection with H3N2 and RSV. The data presented is an average of 3 independent experiments ± standard error.

### Influenza infection inhibits RSV in vivo in a two-wave modality

To test whether influenza infection inhibits RSV in vivo, mice were infected with influenza A/PR8/1934 ("Influenza A") and then infected with RSV (Fig 3A). Influenza A virus was detected in mouse lungs until day 6 post-infection (Fig 3B). In contrast, RSV was not detected at all until day 3 (Fig 3B). From day 3 post Influenza A infection, in parallel to the decline in influenza A virus levels, RSV was detected in the lungs of the mice, reaching a peak on day 5 (Fig 3B). A second wave of inhibition of RSV infection was observed from day 6 until day 12 (Fig 3B), despite absence of detectable influenza virus in the lungs at these time points (Fig 3).

**Fig 3.**
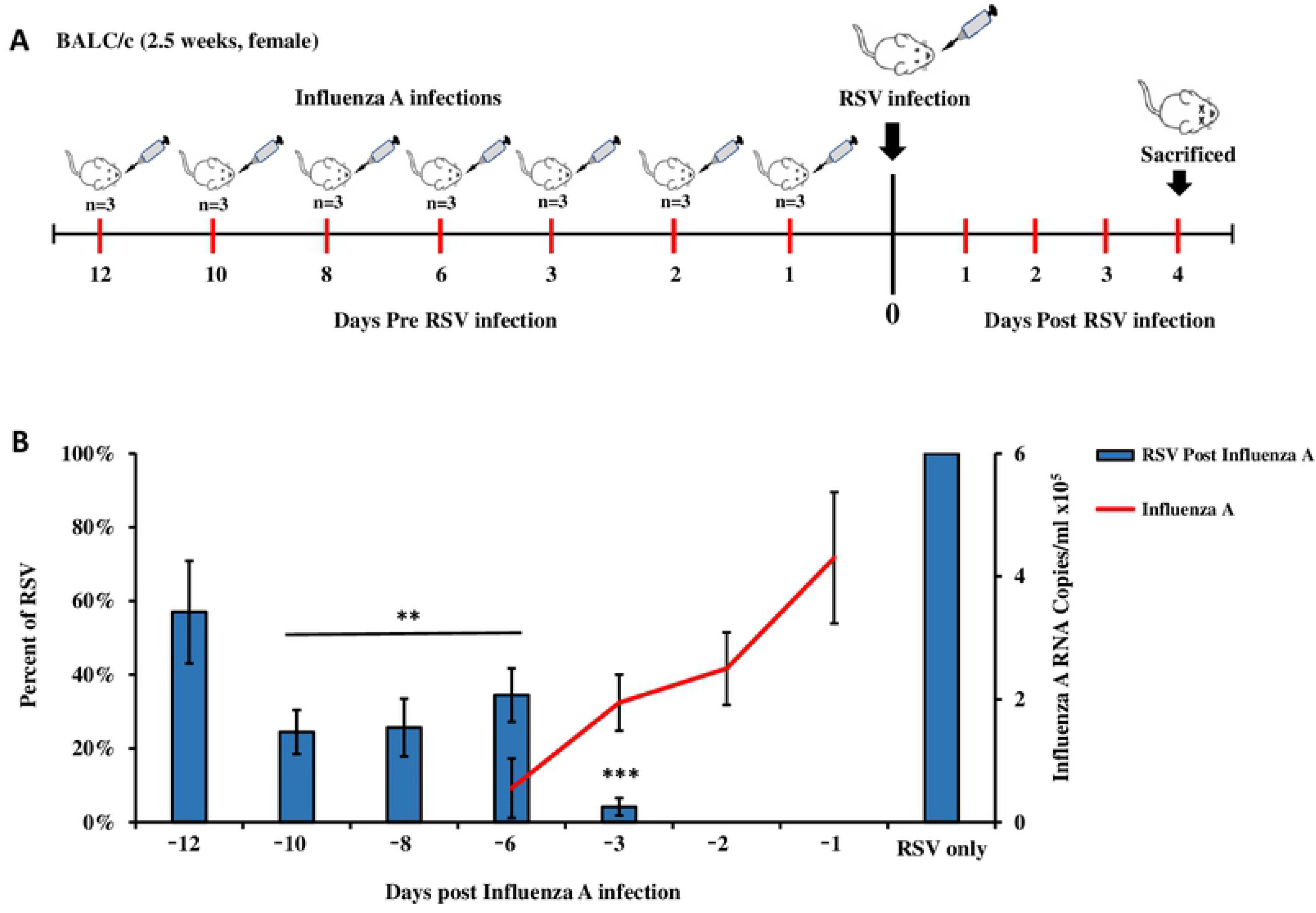
Viral loads in mouse lungs co-infected with influenza (A/PR/8/1934) and RSV. **(A)** Experimental design scheme. 2.5-week-old Balb/c mice intranasally infected with A/PR/8/1934 (4×10^2^ PFU/ml) at predefined time points pre RSV infection (1, 2, 3, 5, 6, 8,10, and 12 days), post A/PR/8/1934 infection the mice were intranasally infected with RSV (6×10^6^ PFU/ml) and then sacrificed four days after RSV infection. mice that were infected with RSV only, served as the control group and were scarified four days later. **(B)** Mouse lungs were homogenized using the Spex centri prep 8000-D Mixer (Mill). RNA was then extracted and the amount of virus was determined by qRT-PCR. Left Y axis represent RSV percent values compare to control group (blue columns), right Y axis represent influenza A/PR/8/1934 RNA copies/ml (Red line).

### Two-wave elevation of anti-viral proteins following influenza infection

To understand the two-wave effect of influenza on RSV, a mass spectrometry analysis was performed on healthy mice and on mice infected with influenza A. A total of 10 proteins demonstrated a two-wave behavior of upregulation (Fig 4) and are known to have antiviral activity (Table 1). Four out the 10 antiviral proteins were IFITs (IFIT1, IFIT2, IFIT3 and IFI44), which were all upregulated on days 1-3 post-infection and then peaked again on day 10 post infection (Fig 5).

**Table 1.** Proteins demonstrated a two-wave behavior of upregulation and are known to have antiviral activity. Table listing the names of 10 proteins shown in Fig 4, which have a known antiviral role, including the IFIT1-3 and IFI44 proteins.

| **Protein names** | **Function** |
| --- | --- |
| Eif2ak2 | Inhibits viral replication via phosphorylation of the alpha subunit of eukaryotic initiation factor. |
| Gbp1 | Promote oxidative killing and deliver antimicrobial peptides to autophagolysosomes. |
| Phf11 | Inhibitor of prototype foamy virus (PFV) replication. |
| Ifi44 | Exhibits an antiviral activity against hepatitis C virus. |
| Ifit1 | Acting as a sensor of viral single- stranded RNAs and inhibiting expression of viral messenger RNAs. |
| Ifit2 | Distinguish between self and non-self mRNAs by the host during viral infection. |
| Ifit3 | IFN-induced antiviral protein which acts as an inhibitor of viral processes, and viral replication. |
| Ifitm3 | Inhibits the entry of viruses to the host cell cytoplasm. |
| Oas3 | dsRNA-activated antiviral enzyme which plays a critical role in cellular innate antiviral response. |
| Zbp1 | Participates in the detection by the host's innate immune system of DNA from viral. |

**Fig 4.**
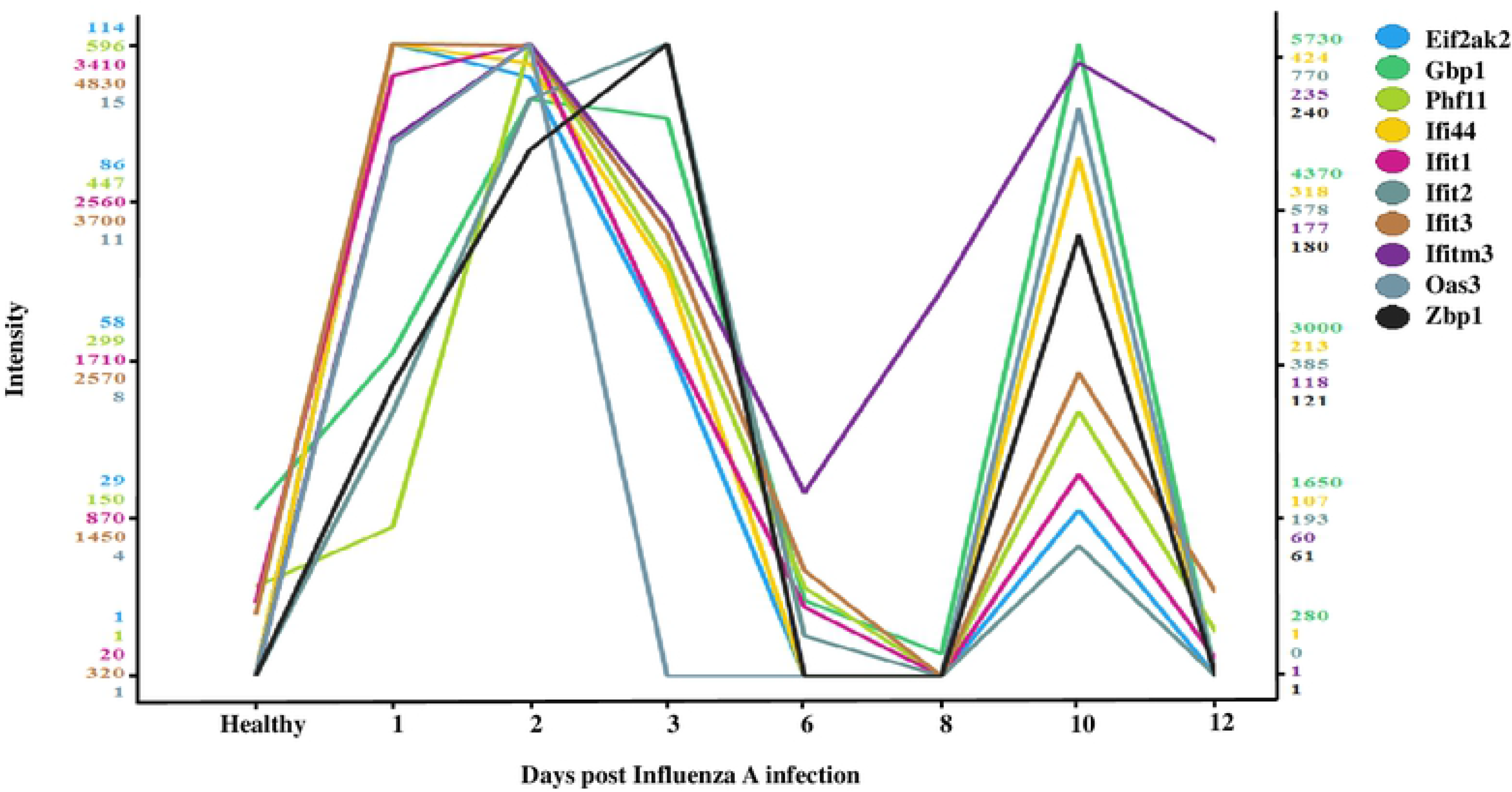
Mass spectrometry analyses of influenza-infected mouse lungs (A/PR/8/1934) versus healthy (non-infected) mouse lungs. Plot of antiviral genes expression levels in influenza A/PR/8/1934 infected cells (2-fold change influenza A/PR/8/1934 vs. healthy) with expression pattern similar to the IFIT1 protein expression pattern (correlation ≥0.6, Tibco Spotfire V.7.7.0. software). The color scale (y axis) for each gene is independent, based on the distribution of the protein’s expression levels across the samples. The samples plotted (x axis) are the healthy, non-infected cells and the post influenza A/PR/8/1934 infection sample from day 1-3, day 6, day 8, day10 and day12.

**Fig 5.**
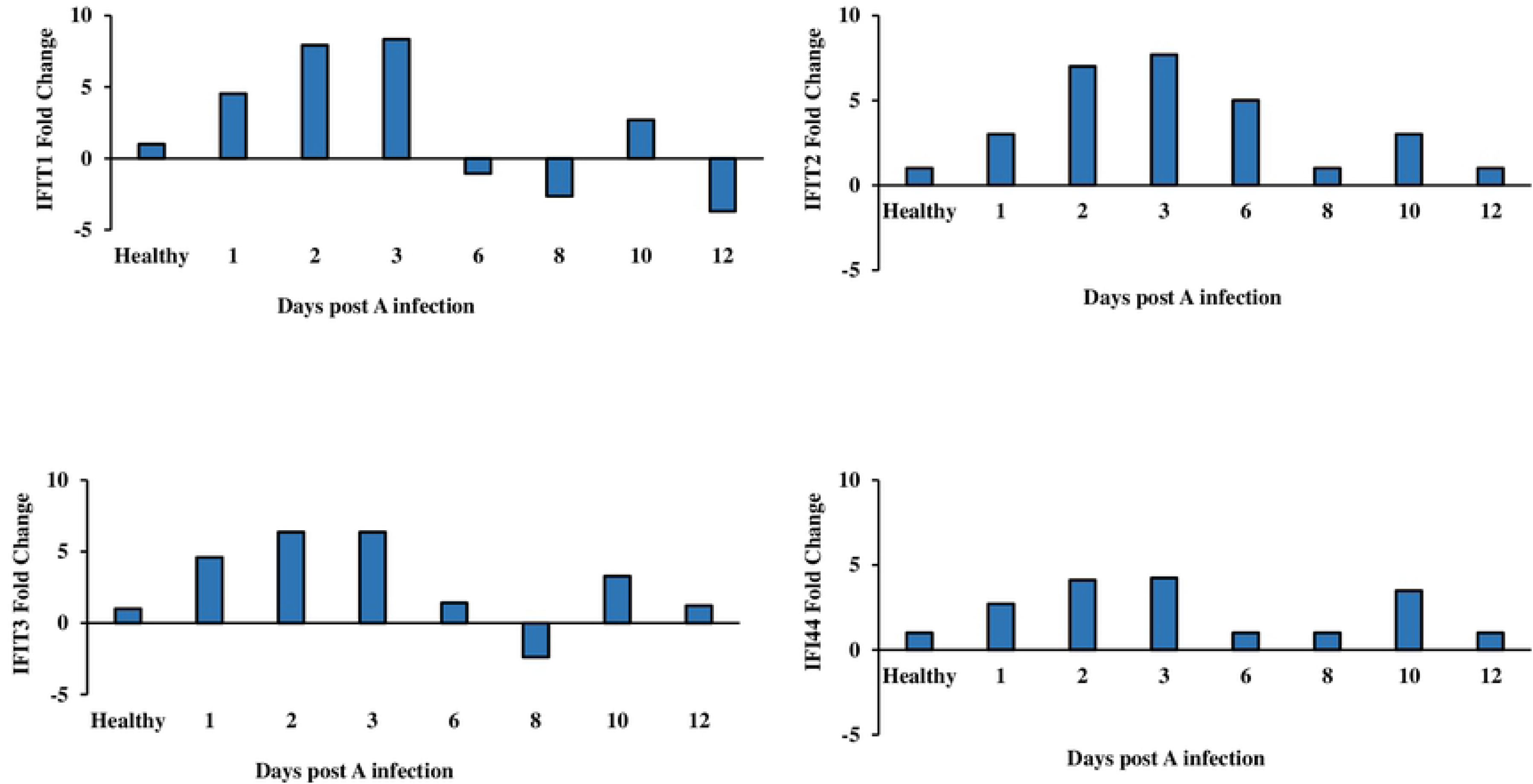
IFIT1-3 and IFI44 protein expression pattern in influenza (A/PR/8/1934)-infected versus healthy mouse lungs. The graphs shown indicates the change in the above proteins expression; upregulated (positive values) or downregulated (negative values) proteins showing a ≥2-fold change in active expression, as analyzed by Tibco Spotfire V.7.7.0.

### IFIT members inhibit RSV infection

To assess the role of IFIT1-3 and IFI44 protein in inhibition of RSV infection, each gene was individually silenced in Hep2 cells. Silencing was confirmed by Western blotting (Fig 6A-D). Silencing of each of the four proteins resulted in increased RSV infection as compared to wild type cells (Fig 6E). In parallel, RSV infection of IFIT1-3 or IFI44-overexpressing HEP2 cells (Fig 7A-D), inhibited the multiplicity rate of RSV starting from 72 h post-infection (p <0.01) (Fig 7E).

**Fig 6.**
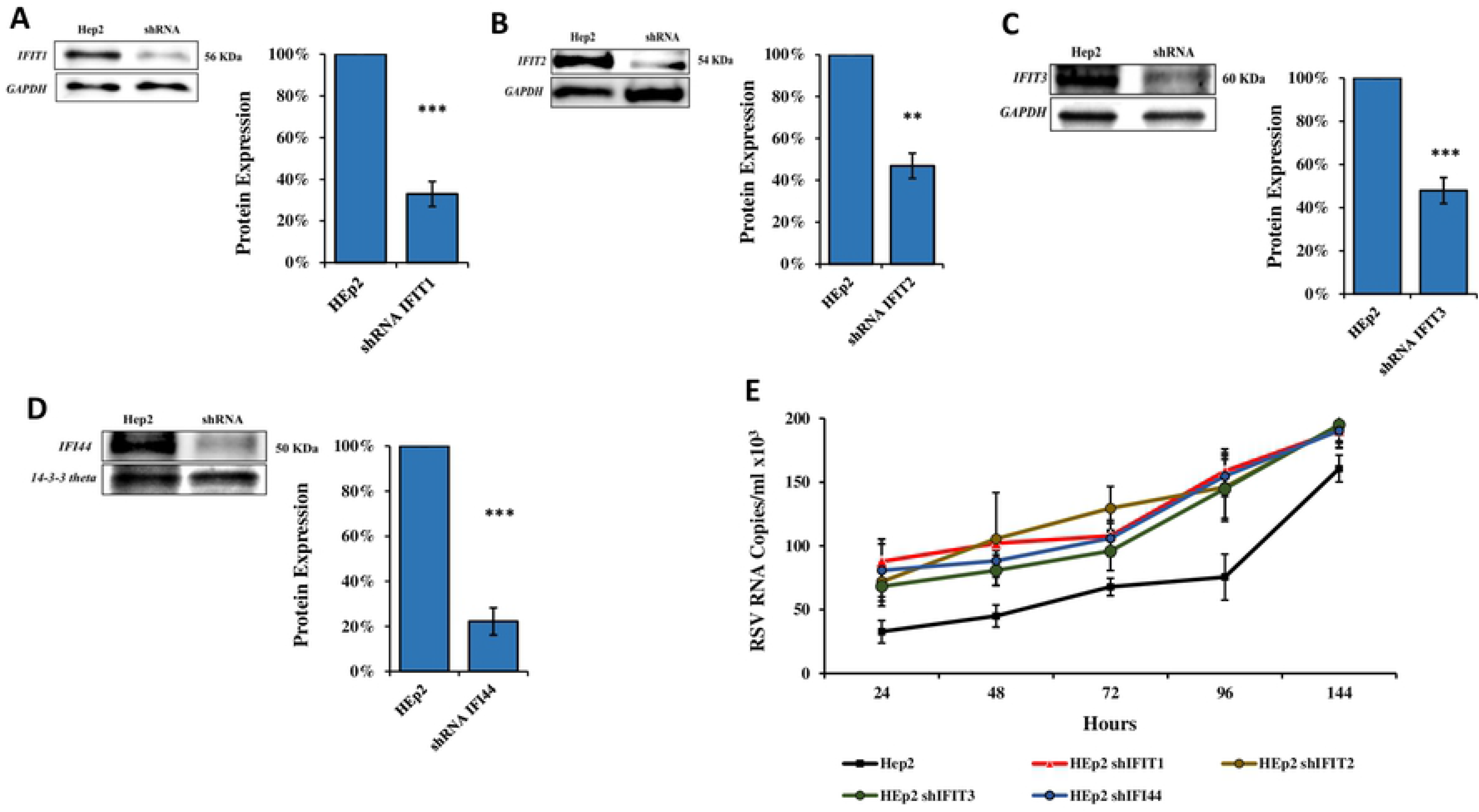
Infection of IFIT1-3 and IFI44 silenced HEp2 cells with RSV. **(A-D)** Lentiviral vectors carrying shRNAs targeting IFIT1, IFIT2, IFIT3 or IFI44 were infected into HEp2 cells. Following selection by puromycin, whole cell extracts were harvested after 5 days of infection. Western blot analysis using antibodies specific for (a) IFIT1, (b) IFIT2, (c) IFIT3 or (d) IFI44. The bars on the graph representing the quantification of IFIT1-3, IFI44. IFIT1-3 and IFI44 expression showed 32.97%, 46.88%, 53.37% and 34.20%, respectively, as compared to untreated control cells. Error bars represent S.E. ** p<0.01, *** p<0.001. **(E**) shIFIT1/shIFIT2/shIFIT3/shIFI44-silenced HEp2 cells (1×10^6^) were infected with RSV (6×10^5^ PFU, 3 h). Supernatant samples were collected at predefined time points (24-144 hours) and RNA was extracted. qRT-PCR analysis was performed to test for viral quantity/presence. The data presented is an average of 3 independent experiments ± standard error.

**Fig 7.**
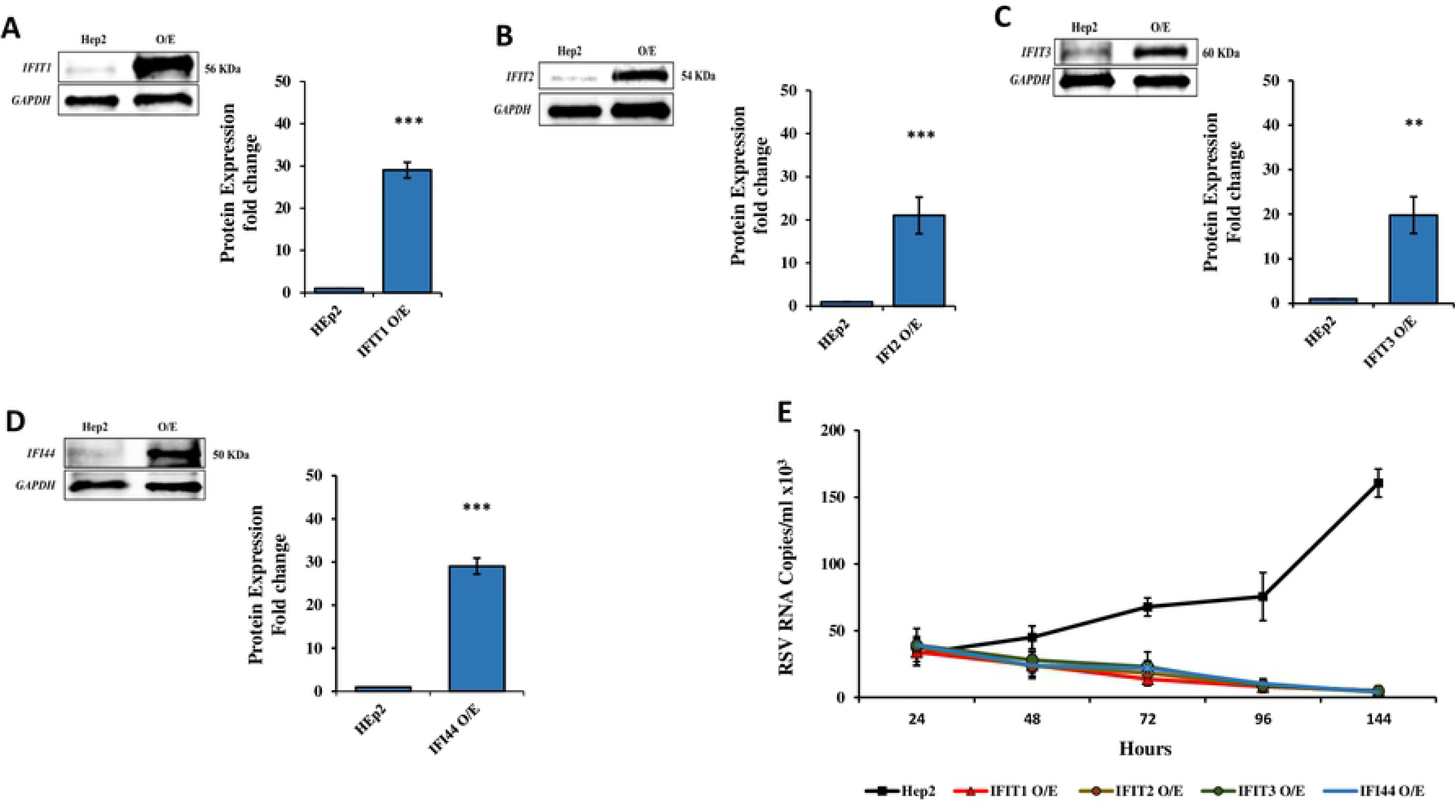
Infection of HEp2 - IFIT1/ IFIT2/ IFIT3/ IF44 Overexpression (O/E) with RSV. **(A-D).** HEp2 cells were transduced with a lentivirus system carrying IFIT1-3 and IFI44 pHAGE-DsRED(−)eGFP(+). Western blots analysis show overexpression of IFIT1-3 and IFI44 expression (29.031%, 21%, 19.8% and 24.20%, respectively, as compared to untreated control cells). Error bars represent S.E. ** p<0.01, *** p<0.001. (**E)** IFIT1/ IFIT2/IFIT3/IF44-overexpressing (O/E) HEp2 cells (1×10^6^) were infected with RSV (6×10^5^ PFU, 3 h). Supernatant was collected at predefined time points thereafter (24-144 hours) and RNA was extracted. qRT-PCR analysis was performed to determine viral load. The data presented is an average of 3 independent replicates ± standard error.

## Discussion

Influenza viruses and RSV are major causes of morbidity and mortality around the world. Generally, in Israel, these viruses are most common during the fall and winter seasons. Although the exact timing and duration of influenza seasons can vary, RSV activity is usually between October and December(8) and influenza activity often begins to increase between December and February(18). The current study noted an unusual overlapping peak of influenza and RSV virus activity in hospitalized patients as well in non-hospitalized patients in the last four winter seasons (2016-20). Furthermore, during these seasons, when RSV appeared together with influenza, the percentages of RSV-infected patients was significantly lower, which was assumed to be due to the early emergence of influenza in these specific seasons. Indeed, a delay in RSV infections was previously reported in winter season 2009-10, most likely due to the emergence of the influenza A/H1N1pdm09 virus in the preceding summer(8). This phenomenon is not unique to Israel, as a similar delay in RSV emergence was reported in various locations in Europe and was also observed with regard to other respiratory viruses, such as seasonal influenza and hMPV(19). In vitro studies support this indication, as it was reported that RSV and influenza share an ecologic niche(20), in which growth of RSV is blocked by competitive infection with influenza(21).

We suspected that influenza and RSV interaction may be affected by the immune system response mechanism. It was reported that continual infection with influenza virus leads to dysregulation of immune responses and hence to a microenvironment supportive of secondary infections (22, 23). In contrast, the present in vitro and in vivo analyses showed that infection with influenza provided resistance to subsequent infection with RSV. More specifically, influenza infection upregulated anti-viral molecules that provided immunity from RSV, and indeed, most co-infections with influenza involve secondary bacterial infections as opposed to other viral infections(24, 25).

The in-vivo experiments demonstrated that infection with influenza virus inhibits the ability of RSV to infect mouse lungs cells. A two-wave modality of inhibition, was noted, with intensified RSV inhibition on days 2-5 and days 8-10 post-infection, with the second wave of RSV inhibition occurring despite the absence of detectable influenza viruses. These waves were paralleled by a two-wave expression pattern of 10 proteins which have known antiviral activity, and four of which belonged to the IFIT family. It was already shown that IFIT1-3 are key molecules in the cells antiviral pathway, where knockout of only one render target cells more vulnerable to viral attack(14). More specifically, IFIT1 acts as a sensor that recognizes non-self RNAs, IFIT2 inhibits their translation or replication at the initiation stage(26), while IFIT3 acts as cofactor that stimulates IFIT1:2 complex stability and activity(27).

The mechanisms underlying the two-wave expression of these 10 proteins are not fully understood. As mentioned, the second wave of protein expression occurred in the absence of influenza virus. Interestingly, the same pattern of immune activity was observed two weeks after inoculation of mice with Sendai virus, when the virus was no longer detectable(28). Acute viral infection induces alteration in the innate immune response, which then drives the development of chronic airway disease, which might then re-activate the innate immune system(29). The presence of a persistent innate immune response on day 10 post-influenza infection suggests that there is ongoing immune stimulation that might be mediated by viruses which are contained in macrophages or other cell types. For example, conventional dendritic cells, which are sites of virus uptake, are capable of activating natural killer T cells at low levels of antigen(30, 31) and may stimulated anti-viral pathways such as the interferon-stimulated genes (ISGs) pathway(32). Future works must therefore focus on the IFIT1-3 and IFI44 proteins, as well as other yet identified proteins that induce chronic innate immune activation after viral infection and on the possibility that viral remnants contained in macrophages or other cell types, drive this process.

Topical prophylactic mucosal application of aminoglycosides has been found to increase host resistance to a broad range of viral infections, including herpes simplex viruses, influenza A virus and Zika viruses. Microarray analysis uncovered a marked upregulation of interferon-stimulated genes (ISGs) including the IFIT protein family following aminoglycoside application, highlighting an unexpected ability of aminoglycoside antibiotics to confer broad antiviral resistance in vivo. However, further understanding the precise mechanism by which aminoglycosides induce ISGs will be useful for the future design of novel anti-viral drug(33).

## Materials and Methods

### Clinical samples

Nasopharyngeal samples were collected from both hospitalized and non-hospitalized patients. Samples from community patients with influenza-like illness (ILI) were collected as part of the seasonal influenza surveillance network, which operates in collaboration with the Israel Center for Disease Control (ICDC) in dozens of clinics in Israel. Samples from hospitalized ILI patients were collected at Sheba Medical Center (SMC). The retrospective analysis of the samples obtained from hospitalized patients was conducted for samples collected in 2009-10 (n=15740). Further analysis was conducted with samples collected from hospitalized patients in the years 2013-14 (n=5098), 2014-15 (n=4653), 2015-16 (n=4982), 2016-17 (n=6098), 2017-18 (n=6579) 2018-19 (n=7280) and 2019-20 (n=7794) and with samples collected from outpatient during the same period (2013-14, n=1755; 2014-15, n=1142; 2015-16, n=1919; 2016-17, n=1284; 2017-18, n=1461; 2018-19, n=1488 and 2019-20, n=2051) except the winter of 2009-10. Samples included were collected between October and April of each year.

### Viral RNA extraction

Viral RNA was extracted from cell supernatants using the MagNA PURE 96 instrument, according to the manufacturer’s instructions. Briefly, nucleic acids were extracted from 500 μl of each sample and eluted in 50μl elution buffer. Real-time polymerase chain reactions (qPCR) were performed using the Ambion Ag-Path master mix (Life Technologies, USA) and the ABI 7500 instrument, to test for the presence of influenza viruses as previously described(8, 34) Determination of human RSV, was performed using qPCR, as previously described(8).

### Co-infection of Hep2 cells with RSV and influenza A/H3N2

Human cervical carcinoma (HEp2; ATTC) cells (1×10^6^), which are susceptible to both influenza virus and RSV^19^, were grown in 6-well plates (Nunc™) in 2 ml 10% fetal calf serum (FCS)-enriched MEM-Eagle, Earle’s salts (Biological Industries, Israel), and incubated overnight (37°C, 5% CO_2_). The cells were washed with phosphate buffered solution (PBS) and infected with RSV (6×10^5^ PFU). After 3 h, the cells were washed with PBS then infected with influenza A/H3N2 (6×10^5^ PFU) for 3 h. Following washing with PBS, the cells were incubated (37°C, 5% CO_2_) in 3 ml 2% FCS-enriched MEM-Eagle, Earle’s salts. Supernatant was collected at predefined time points (24, 48, 72, 96 and144 hours) and RNA was extracted. qPCR was performed to quantitate the level of the virus at each time point.

### Co-infection of mice with influenza A/PR/8/1934 and RSV

2.5-week-old *Balb/c* mice were anesthetized with isoflurane (Piramal Critical Care, Inhalation liquid) and intranasally infected with influenza A/PR/8/1934 (“influenza PR8”) (4×10^2^ PFU/ml). The mice were split into groups according the days of post influenza A/PR/8/1934 infection (1, 2, 3, 5, 6, 8, 10, 12), three mice from each group were subsequently infected with RSV (6×10^6^ PFU/ml) for 4 days and then sacrificed. The presence of the viruses was tested by grinding the lungs of the mice in 1ml DMEM (Biological Industries, Israel) using the Spex centri prep 8000-D Mixer Mill homogenizer (BENCHMARK SCIENTIFIC), for 10 min, followed by RNA extraction and qRT-PCR, as described(8) and was compared to control mice (mice whom infected with a single virus; RSV only).

### Mass spectrometry

Infected mouse lung samples were homogenized in 1 ml DMEM (Biological Industries, Israel) with protease inhibitor “cOmplete” ULTRA Tablets, Mini, EASYpack (ROCH) 1 tab\1 ml DMEM and analyzed by mass spectrometry at the Smoler Proteomics Center, Technion, Haifa Israel(35).

### Genes silencing of IFIT1, IFIT2, IFIT3 and IFI44

Lentiviral particles of *sh*RNA specific for human IFIT1, IFIT2, IFIT3 and IFI44 as well as non-targeting *sh*RNA, were purchased from Sigma-Aldrich. Transduction into HEp2 cells was performed according to the manufacturer’s instructions. HEp2 clones were selected with puromycin 6 μg/ml (Sigma-Aldrich) and viral RNA was detected by RT-PCR performed using the following primers:

IFIT1
forward, 5’-TCTCAGAGGAGCCTGGCTAA-3’ reverse, 5’-TCAGGCATTTCATCGTCATC-3’
IFIT2
forward, 5’-CGAACAGCTGAGAATTGCAC-3’ reverse, 5’-TGCACATTGTGGCTTTGAAT-3’
IFIT3
forward, 5’-CGGAACAGCAGAGACACAGA-3’ reverse, 5’-CTGCCTCGTTGTTACCATCT-3’
IFI44
forward, 5’-AGCTGGGAAGTCCAGCTTTT-3’ reverse, 5’-CCCCAGTGAGTCACACAGAA-3’
The silenced clones named: “HEp2 *shIFIT1,* HEp2 *shIFIT2,* HEp2 *shIFIT3* and HEp2*shIFI44*”

### Gene overexpression of IFIT1, IFIT2, IFIT3 and IFI44

Lentiviral vectors were produced in 293T cells using the TransIT-LT1 transfection reagent (Mirus, ZOTAL), in a transient three-plasmid transfection protocol according to the manufacturer’s instructions. The expression vector used was pHAGE-DsRED(−) eGFP(+). Primers designed to create the insert were as follows:

IFIT1
forward, 5’-TTTTCTCGAGGCCGCCACCATGAGTACAAATGGTGATGATC-3’ reverse, 5’-AAAGCTAGCCTAAGGACCTTGTCTCACAGAGTTC-3’
IFIT2
forward, 5’-TTTTCTCGAGGCCGCCACCATGAGTGAGAACAATAAGAA-3’ reverse, 5’-AAAGCTAGCTCATTCCCCATTCCAGCTTGATGCT-3’
IFIT3
forward, 5’-TTTTCTCGAGGCCGCCACCATGAGTGAGGTCACCAAGAATTC-3’ reverse, 5’-AAAGCTAGCTCAGTTCAGTTGCTCTGAGTTAG-3’
IFI44
forward, 5’-TTTTCTCGAGGCCGCCACCATGGCAGTGACAACTCGTTTGAC-3’
reverse, 5’-CCCGCTAGCCTACCCGCTAGCCTATTTTTTTCCTTGTGCACAGTTGATAATTTCCTCCCTTAGATTC-3’
The overexpression clones named: “IFIT1 O/E, IFIT2 O/E, IFIT3 O/E and IFI44 O/E”

### Western blot

Whole-cell extracts were eluted using Pierce™ RIPA Buffer (Thermo SCIENTIFIC, USA), according to the manufacturer’s protocols, fractionated by SDS-PAGE 4-20% (Mini-PROTEAN TGX Stain-free Gels, BIO-RAD, USA) and transferred to a 0.2μm nitrocellulose membrane (Trans-Blot Turbo Transfer Pack, BIO-RAD, USA) using a Turbo® Transfer System, according to the manufacturer’s protocols (BIO-RAD, USA). After incubation with InstantBlock buffer (Gene Bio-Application L.T.D, Israel) according to the manufacturer’s protocols, the membrane was washed once with TBST (10 mM Tris, pH 8.0, 150 mM NaCl, 1% Tween 20) and incubated with polyclonal antibodies (Thermo SCIENTIFIC, USA) specific for IFIT1 (1:1000), IFIT2 (1:1000), IFIT3 (1:1000), IFI44 (1:1000) or GAPDH (1:5000) for 1-3 h, at 4°C. Membranes were washed three times and incubated with a 1:10,000 dilution of horseradish peroxidase-conjugated anti-rabbit antibodies ((Sigma-Aldrich) for 1-2 h, at room temperature. Blots were washed with TBST and developed with the Clarity™ Western ECL substrate (BIO-RAD, USA), according to the manufacturer’s protocols.

### Infection of IFIT-silenced/overexpressed HEp2 cells with RSV

IFIT-silenced/overexpressed HEp2 (1×106) cells were grown overnight in 6-well plates (NuncTM) in 10% FCS-enriched EMEM (37°C, 5% CO2). The cells were washed with 1 ml PBS and infected with RSV (6×10^5^ PFU). After 3 h, the cells were washed with 1 ml PBS, and incubated in 2% FCS-enriched MEM-Eagle, Earle’s salts (37°C, 5% CO_2_), for 5 days. Supernatant were taken to test for the presence of human RSV and was performed by using qPCR, as previously described(8).

### Statistical analysis

T-test was applied to evaluate the differences in percent positivity between the compared groups. A p value < 0.05 was considered statistically significant. All analyses were performed using IBM® SPSS® Statistics software (Version 23) and Excel software (Microsoft®).

**This work was performed in partial fulfillment of the requirements for the Ph.D degree of Yaron Drori, Sackler Faculty of Medicine, Tel Aviv University.**

